# OAS3 and RNase L integrate into higher-order antiviral condensates

**DOI:** 10.1101/2023.05.30.542827

**Authors:** Renee Cusic, James M. Burke

## Abstract

Oligoadenylate synthetase 3 (OAS3) and Ribonuclease L (RNase L) are components of a mammalian RNA decay pathway that is critical for combating viral infection. Nevertheless, the subcellular localization of OAS3 and RNase L during the antiviral response has remained elusive. Herein, we show that upon activation in response to double-stranded RNA or viral infection, OAS3 and RNase L integrate into higher-order cytoplasmic assemblies distinct from stress granules (SGs), RNase L-induced bodies (RLBs), or processing bodies (P-bodies). We identified these assemblies as double-stranded RNA-induced foci (dRIF), which also contain activated protein kinase R (PKR). We observed incorporation of RNase L, OAS3, and PKR in dRIFs during Zika virus and dengue virus infection. Our data support that condensation of dsRNA, OAS3 and RNase L at dRIFs promotes the homodimerization and oligomerization of RNase L required for its rapid activation, demonstrating a fundamental role for biological condensates in activating antiviral signaling.

## INTRODUCTION

Ribonuclease L (RNase L) is a ubiquitously expressed endoribonuclease in mammals that combats viral infection and tumorigenesis (Silverman 2007; Chakrabarti et al., 2011). While latent under homeostatic conditions, RNase L is activated when cytoplasmic levels of double stranded RNA (dsRNA) increase, which is common during viral infection or dysregulation of endogenous RNAs (Floyd-Smith et al., 1981; Li et al., 2016; Li et al., 2017). RNase L is not directly activated by dsRNA. Instead, binding of dsRNA by oligoadenylate synthetase (OAS) proteins triggers the generation 2’-5’ oligo(A) (2-5A). The binding of 2-5A at the N-terminal domain of RNase L allows for homodimerization of RNase L, which activates its C-terminal RNase domain. Activated RNase L homodimers then oligomerize into the fully activated form of RNase L (Han et al., 2012). Once activated, RNase L cleaves ssRNA regions at UNN motifs in most types of RNAs, including cellular tRNAs, rRNAs, and mRNAs and viral genomes/mRNAs (Wreschner, McCauley et al., 1981; Silverman et al., 1983; Wreschner, James et al., 1981; Li et al., 1998; Khabar et al., 2003; Andersen et al., 2009; Brennan-Laun et al., 2014; Donovan et al., 2017; Rath et al., 2019; Burke et al., 2019).

The degradation of cellular RNA within single cells is rapid and robust upon activation of RNase L, whereby most cellular mRNAs are completely degraded within one-hour post activation of RNase L (Burke et al., 2019; Cusic et al., 2023). The bulk degradation of mRNAs regulates several cellular processes. First, it reprograms the transcriptome to an antiviral state because immediate early antiviral mRNAs (i.e., type I and III interferons) evade RNase L mediated mRNA decay. Second, it inhibits the assembly of stress granules (SGs) (Burke et al., 2019), which are large cytoplasmic ribonucleoprotein (RNP) complexes that concentrate non translating mRNAs and RNA-binding proteins such as G3BP1 and poly(A)-binding protein (PABP) during stress-induced translational repression (Buchan and Parker, 2009). Third, it induces the assembly of small RNP complexes termed RNase L-induced bodies (RLBs), which are similar to SGs in that they contain G3BP1, PABP, and poly(A)+ RNA, but are distinct from SGs in their morphology, overall composition, and biogenesis (Burke et al., 2020; Burke, 2022). Fourth, it causes RNA-binding proteins such as PABP to translocate from the cytoplasm to the nucleus, which leads to nuclear RNA processing and mRNA export defects (Burke, Ghilcrest et al., 2021; Burke et al., 2022).

Despite the importance and numerous cellular processes regulated by the OAS/RNase L pathway during the antiviral response, limited information is known about the subcellular localization of OAS proteins and RNase L. While RNase L and OAS proteins are generally thought to reside in the cytoplasm, RNase L has been reported to localize to the nucleus in a cell cycle-dependent manner (Al-Ahmadi et al., 2009). In addition, both OAS proteins and RNase L have been reported to localize to stress granules (Reineke and Lloyd, 2015; Manivannan et al., 2020; Paget et al., 2023). Lastly, whether OAS and RNase L localize to RNase L-induced bodies (RLBs) has not been examined.

Herein, we comprehensively examine the subcellular localization of OAS3 and RNase L using live-cell fluorescence microscopy during the innate immune response to dsRNA or viral infection. We observed that in response to dsRNA, Zika virus infection, and dengue virus infection, OAS3 and RNase L integrate into higher-order antiviral signaling complexes. We identified these structures as double-stranded RNA-induced foci (dRIF). Contrary to previous reports, we observed that neither OAS3 nor RNase L localize to G3BP1 RNP complexes, including SGs and RLBs. Our data support a model whereby concentration of dsRNA, OAS3 and RNase L in dRIFs promotes homodimerization and oligomerization RNase L, leading to rapid activation of RNase L in response to viral infection. These findings demonstrate that biological condensates play a key role regulating antiviral signaling pathways.

## RESULTS

### GFP-tagged RNase L mimics endogenous RNase L latency and activation

The sub-cellular localization of RNase L has remained enigmatic. To address this, we initially performed immunofluorescence assays for RNase L. However, two commercial antibodies failed to provide RNase L-specific immunofluorescence (Fig. S1A). Therefore, we introduced stable expression of RNase L with an N-terminal GFP tag (GFP-RNase L) via lentivirus transduction in RNase L-KO A549 cells (Fig. 1A,B). Because the N-terminal GFP tag is near the regulatory domain of RNase L (2-5A binding) (Fig. 1A), we examined whether it altered the ability of RNase L to remain latent and/or properly activate.

**Figure 1.**
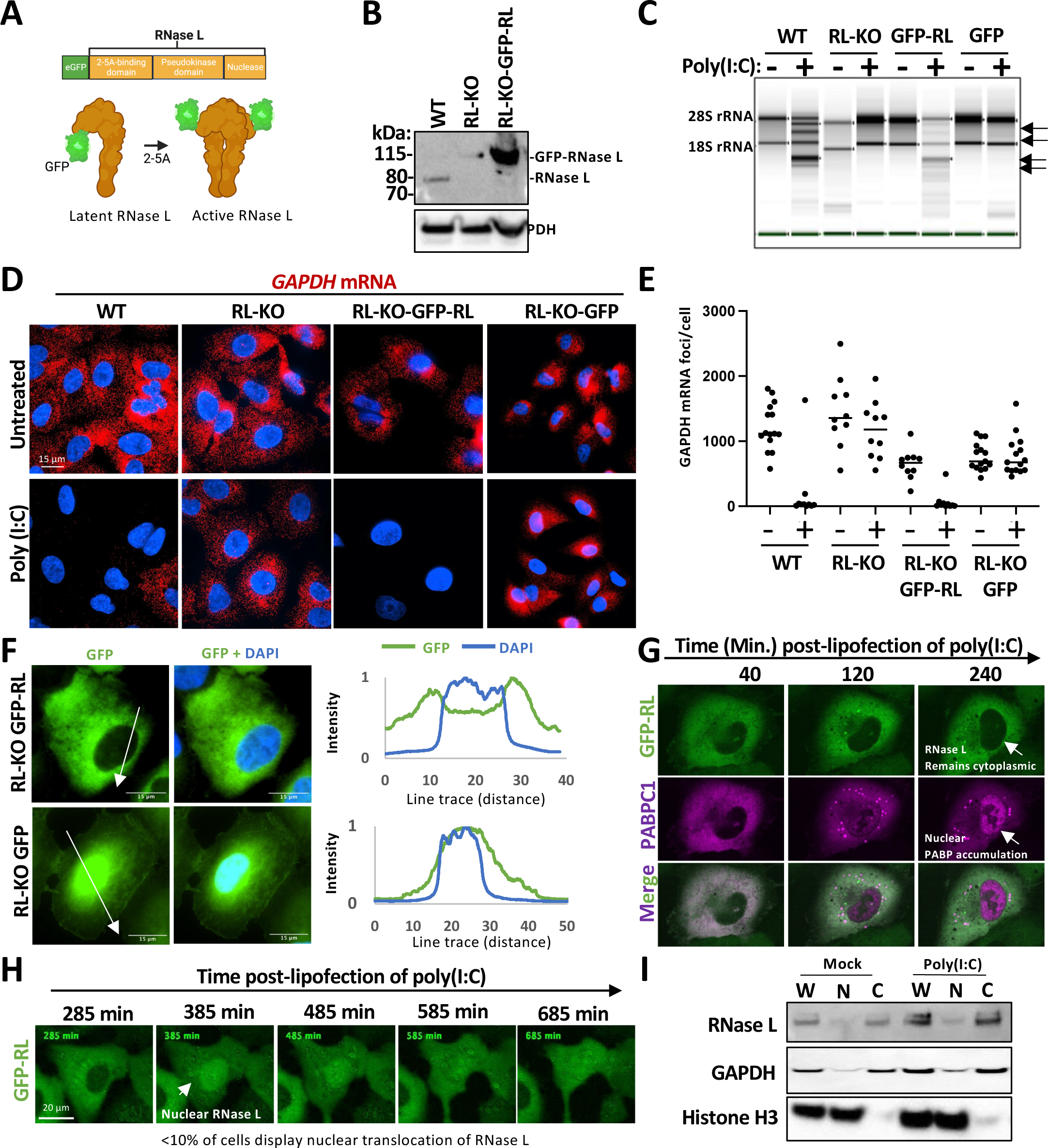
N-terminally GFP tagged RNase L is functional and localizes to the cytoplasm. (A) Model showing GFP tag relative to RNase L domains. (B) Western blotting analyses in wild-type (WT), RNase L-KO (RL-KO), and RL-KO cells stably expressing GFP-RNase L (GFP-RL) A549 cell lines. (C) TAPE station analysis for RNase-dependent cleavage of ribosomal RNA (arrows) following lipofection with or without poly(I:C). (D) smFISH staining for *GAPDH* mRNA. (E) Quantification of *GAPDH* mRNA foci/ cell as represented in (D). 15 cells from each cell line were analyzed using 2-3 fields of view. (F) Fluorescence microscopy in RL-KO cells expressing with GFP-RNase L or the GFP-only control. Line traces show normalized intensity of DAPI and GFP staining. (G) Live cell imaging of RL-KO cells stably expressing GFP-RNase L and mRuby2-PABPC1 transfected with poly(I:C). (H) Live cell imaging of RL-KO cells expressing GFP-RNase L and transfected with poly(I:C). (I) Western blot analysis of crude nuclear/cytoplasmic fractionations.

Importantly, we observed that GFP-RNase L is latent under homeostatic conditions because RNase L-KO cells expressing GFP-RNase L did not display RNase L-dependent ribosomal RNA (rRNA) cleavage fragments (Fig. 1C) or reduced *GAPDH* mRNA levels (Fig. 1D,E) in comparison to the GFP-only control. Following lipofection poly(I:C) (a synthetic dsRNA that leads to RNase L activation), GFP-RNase L rescued cleavage of rRNA (Fig. 1C) and degraded *GAPDH* mRNA (Fig. 1D,E) in RL-KO cells. The GFP-only control did not rescue RNA decay in RL-KO cells (Fig. 1C,D,E). The magnitude by which GFP-RNase L degraded rRNA and mRNA was comparable to those observed in WT cells (Fig. 1C,D,E). These data demonstrate that GFP-RNase L functions similarly to endogenous RNase L, whereby GFP-RNase L is latent under homeostatic conditions but activated in response to double-stranded RNA.

### RNase L is predominately localized to the cytoplasm

We next sought to determine the cellular localization of RNase L via fluorescence microscopy in cells expressing GFP-RNase L or the GFP-only control. We observed that GFP-RNase L diffusely localized to the cytoplasm and was depleted from the nucleus (Fig. 1F). In contrast, the GFP-only control concentrated in the nucleus (Fig. 1F). Importantly, these data indicate that the localization of GFP signal to the cytoplasm is dictated by RNase L in cells expressing GFP-RNase L. Thus, GFP-RNase L fluorescence is a reliable means by which to assess RNase L localization.

Although we observed RNase L was predominately localized to the cytoplasm (Fig. 1F), which is consistent with our previous fractionation studies (Burke et al., 2022), we wanted to determine if RNase L localizes to the nucleus during various cellular states. Because RNase L was reported to localize to the nucleus in a cell cycle-dependent manner, whereby RNase L concentrates in the nucleus in sub-confluent cells but is predominantly cytoplasmic in confluent cells (Al-Ahmadi et al., 2009), we first analyzed RNase L localization during the cell cycle via live-cell imaging. In contrast to Al-Ahmadi et al., we did not observe RNase L concentrate in the nucleus in sub-confluent cells (Fig. S2A). Moreover, RNase L remained cytoplasmic as cells became confluent (Fig. S2A). Lastly, RNase L remained predominantly localized to the cytoplasm throughout multiple cell divisions (Fig. S2B). These data indicate that RNase L localizes to the cytoplasm during homeostatic conditions independent of the cell cycle.

Activation of RNase L-mediated mRNA decay results in the translocation of serval RNA-binding proteins, such as poly(A)-binding protein C1 (PABPC1), from the cytoplasm to the nucleus (Burke et al., 2022). Thus, we reasoned that RNase L could similarly translocate to the nucleus upon activation. However, live-cell imaging of GFP-RNase L cells co-expressing mRuby2-PABPC1 demonstrated that RNase L typically remains localized to the cytoplasm while PABPC1 translocated to the nucleus following poly(I:C) lipofection (Fig. 1G; Movie 1). Thus, RNase L typically remains localized to the cytoplasm upon activation.

We did observe a small percentage (<10%) of cells in which RNase L translocated to the nucleus (Movie 2) (Fig. 1H). However, crude cytoplasmic-nuclear fractionation showed only a modest increase of RNase L in the nucleus following poly(I:C) lipofection (Fig. 1I), indicating that nuclear RNase L localization is rare, transient, or associated with cell death. In support of the latter, GFP-RNase L translocation to the nucleus coincided with a marked cytopathic effect (Fig. 1H and Movie 2). Combined, these data support that RNase L is predominantly localized in the cytoplasm when both latent and active.

### RNase L assembles into granules in response to double-stranded RNA

An important observation we made is that cells expressing GFP-RNase L form large GFP-positive granules following poly(I:C) lipofection (Fig. 2A,B), which we term RNase L granules. Live-cell imaging revealed that RNase L granules are transient, often assembling and disassembling within 20 minutes at early times (0-4 hrs.) following poly(I:C) (Fig. 2C) (Movie 3). Thus, at these early times post poly(I:C) lipofection, only a small fraction (∼10%) of cells display RNase L granules at a given time (Fig. 2D,E). However, RNase L granules became more abundant and/or stable at late times (8-20 hrs.) post-lipofection of poly(I:C), with ∼50% of cells containing multiple RNase L granules (Fig. 2D,E and Movies 4 and 5).

**Figure 2.**
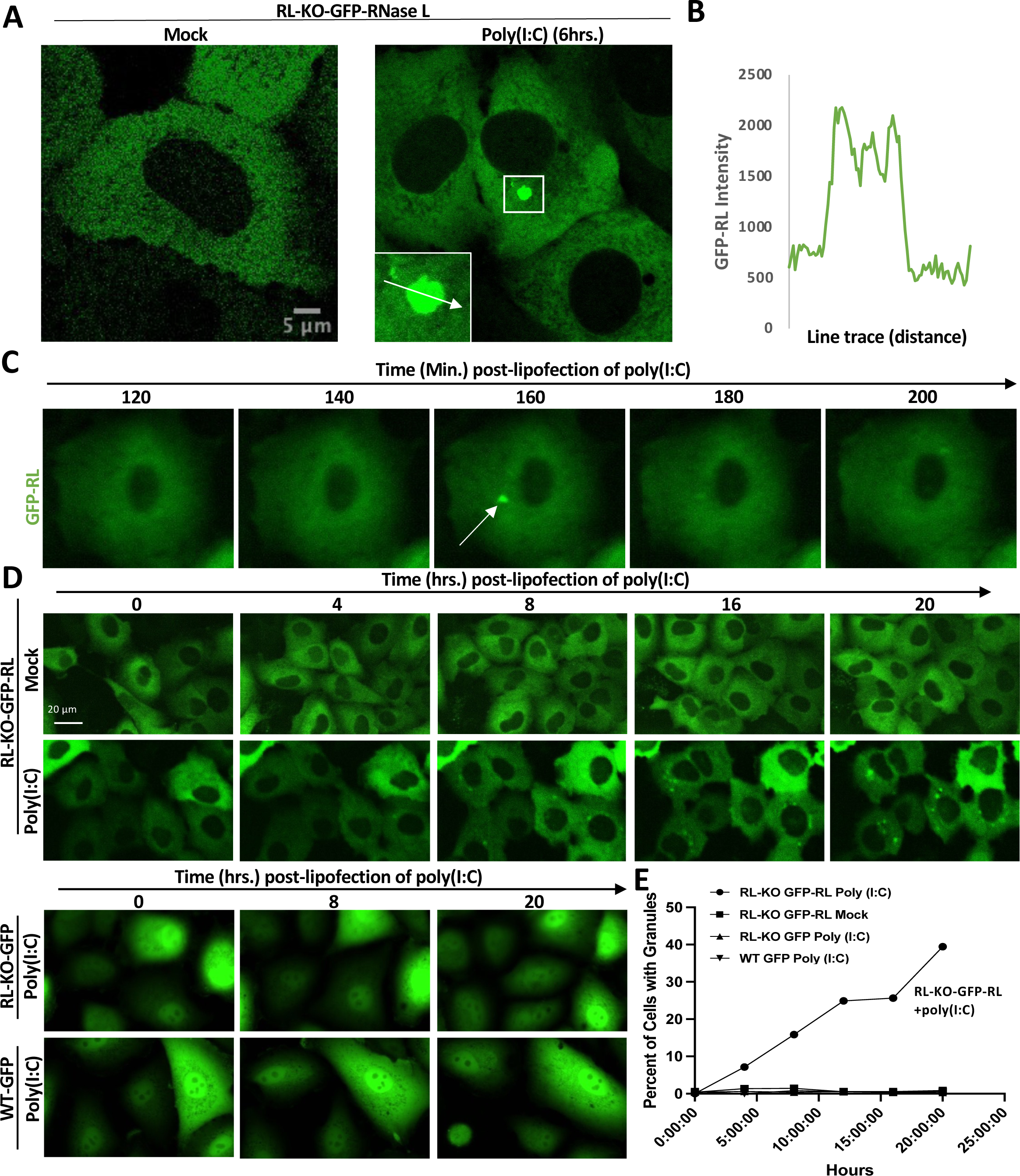
RNase L integrates into high-order assemblies upon activation. (A) Fluorescence microscopy of RL-KO expressing GFP-RNase L displaying a higher-order assembly following poly(I:C) transfection. B) Line trace of granule from (A). (C) Live cell imaging of RL-KO A549 cells expressing GFP-RNase L imaged every 20 minutes following lipofection poly(I:C). (D) Live cell imaging of indicated cell lines expressing either GFP-RNase L or GFP-only following poly(I:C) lipofection. (E) Quantification of the percent of cells with at least one GFP-positive granule as represented in (D).

The GFP-positive granules in GFP-RNase L cells are dependent on GFP fusion to RNase L based on two observations. First, we did not observe GFP-positive foci in RNase L-KO cells expressing the GFP-only control following lipofection of poly(I:C) (Fig. 2D,E, RL-KO-GFP). Second, we also considered the possibility that RNase L activation could promote the assembly of GFP into granules independently of fusion to RNase L. Therefore, we introduced expression the GFP-only control in WT A549 cells (WT-GFP), which robustly activate RNase L (Fig. 1C,D,E). However, we did not observe GFP-positive granules in WT-GFP cells following activation of RNase L via poly(I:C) lipofection (Fig. 2D,E). These data indicate that GFP-positive granules observed in cells expressing GFP-RNase L are driven by RNase L.

### RNase L-positive granules are OAS3-dependent and concentrate activated RNase L

The assembly of RNase L granules following poly(I:C) lipofection suggests that their formation correlates with RNase L activation. To further test this hypothesis, we first asked whether RNase L granule assembly correlated with the assembly of RNase L-induced bodies (RLBs) and the translocation of RNA-binding proteins to the nucleus, which both occur immediately following RNase L-mediated mRNA decay (Burke et al., 2019; Burke et al., 2020). To do this, we performed live-cell imaging in GFP-RNase L cells that co-express mRuby2-PABPC1, which localizes to RLBs and translocates to the nucleus following RNase L activation (Burke et al., 2020). Strikingly, we observed that the assembly of RNase L granules often directly preceded or coincided with RLB assembly and PABPC1 translocation to the nucleus (Fig. 3A,B and Movie 6). This suggests that RNase L granules assemble upon activation of RNase L-mediated mRNA decay.

**Figure 3.**
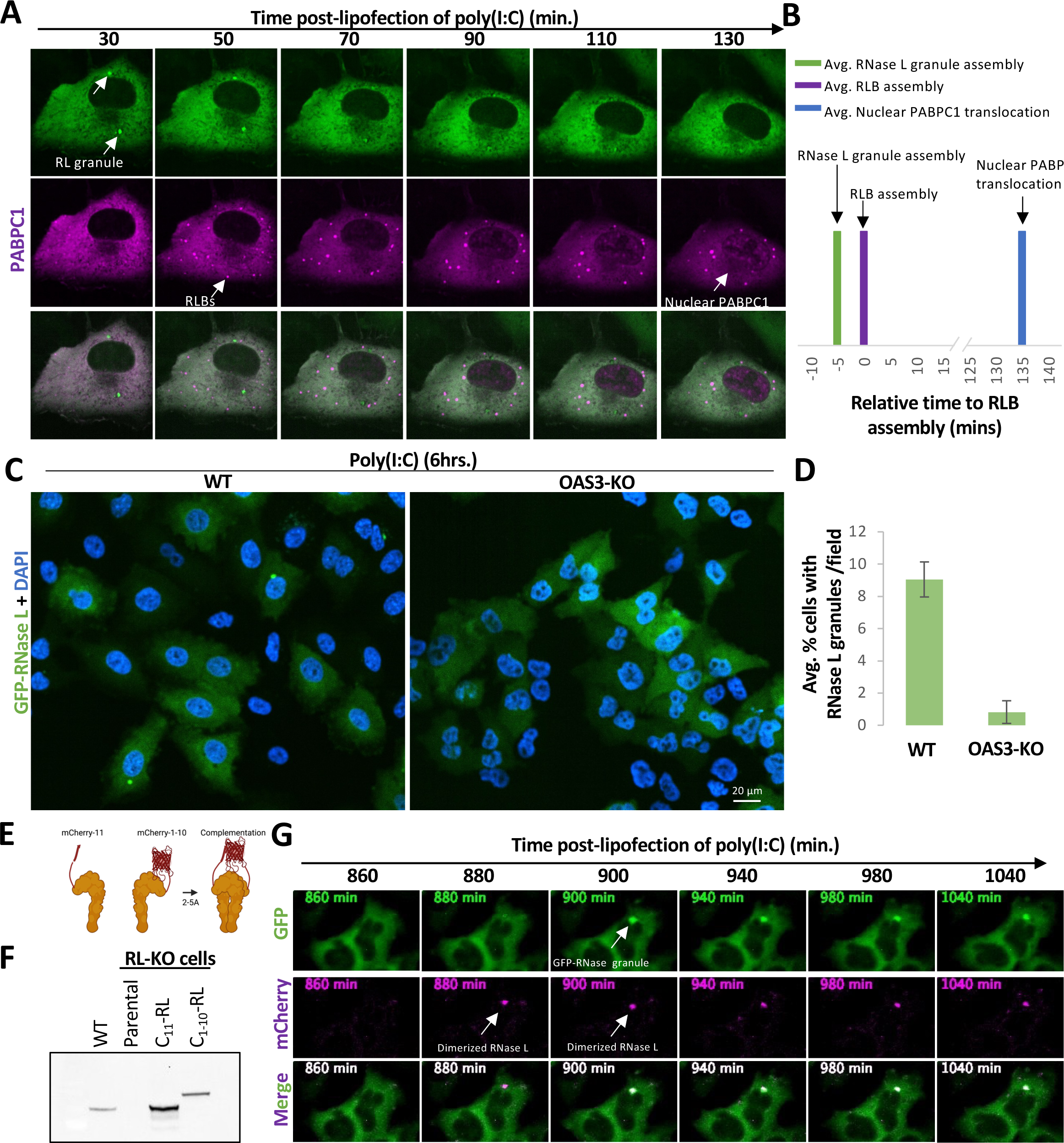
RNase L granules are OAS-dependent and concentrate activated RNase L. (A) Live cell imaging of RL-KO A549 cells co-expressing GFP-RNase L and mRuby2-PABPC1 following poly(I:C) lipofection. (B) Bars represent the average time that RNase L granules (green) and maximum nuclear PABPC1 intensity (blue) occur relative to the time that RNase L-induced bodies (RLBs) form (magenta). (C) Fluorescence microscopy of RNase L-KO or OAS3-KO cells expressing GFP-RNase L six hours post-lipofection of poly(I:C). (D) Average percent +/-SD of cells with GFP-RNase-L assemblies as represented in (C). (E) Model describing split mCherry tagged RNase L. (F) Western blot analysis confirming expression of mCherry_11_-RNase L (C_11_-RL) or mCherry_1-10_-RL (C_1-10_-RL). (G) Live cell imaging of GFP-RNase L and split mCherry-RNase L expressing cells following lipofection of poly(I:C).

The observation that RNase L granules assemble following RNase L activation indicates that RNase L granules are a direct consequence of RNase L activation. Thus, we hypothesized RNase L granules would be dependent on OAS3 for their formation because OAS3 produces 2-5A required for RNase L activation in response to dsRNA (Li et al., 2016). Consistent with OAS3 being required for RNase L activation, most OAS3-KO cells formed stress granules instead of RLBs and lacked nuclear PABPC1 staining following poly(I:C) lipofection (Fig. S3).

Importantly, we observed a 9-fold reduction in the number for cells that formed RNase L granules in OAS3-KO cells expressing GFP-RNase L following poly(I:C) lipofection (Fig. 3C,D). The small percent of OAS3-KO cells expressing GFP-RNase L that formed GFP-positive granules following poly(I:C) (Fig. 3D) typically displayed markers of RNase L activation (nuclear PABP translocation and RLB assembly) (Fig. S3), suggesting these OAS3-KO cells likely activated RNase L via OAS1 or OAS2. Nevertheless, these data support that RNase L granule assembly depends on activation of RNase L.

Because RNase L granules are dependent on RNase L activation, this suggests that they concentrate dimerized/oligomerized (activated) RNase L. To test this, we generated bimolecular fluorescence complementation (BiFC) RNase L reporter cells that co-express RNase L tagged with mCherry_11_, mCherry_1-10_ (Fig. 3E,F) (Feng et al., 2019), which we reasoned would complement upon RNase L dimerization and/or oligomerization. We then performed live cell imaging following poly(I:C) transfection. Remarkably, we observed mCherry-positive granule assembly typically precede or coincide with GFP-RNase L granule assembly (Fig. 3G and Movie 7), indicating that RNase L granules contain dimerized/oligomerized (activated) RNase L. Combined, these data indicate that RNase L granules are dependent on RNase L activation for their formation and concentrate activated (dimerized/oligomerized) RNase L.

### RNase L does not localize to stress granules, p-bodies, or RNase L-induced bodies

We next wanted to determine if RNase L granules are distinct from other cytoplasmic granules that assemble during the antiviral response. We first examined if RNase L localizes to RNase L-induced bodies (RLBs) or processing-bodies (P-bodies), which are both present upon RNase L activation (Burke et al., 2020). Immunofluorescence assays for G3BP1 (RLB marker) and DCP1b (P-body marker) in cells expressing GFP-RNase L following poly(I:C) lipofection demonstrated that RNase L granules are distinct from both RLBs and P-bodies (Fig. 4A). Notably, RNase L did not enrich in either RLBs or P-bodies following poly(I:C) lipofection (Fig. 4A,B), indicating that RNase L does not localize to RLBs or P-bodies when activated.

**Figure 4.**
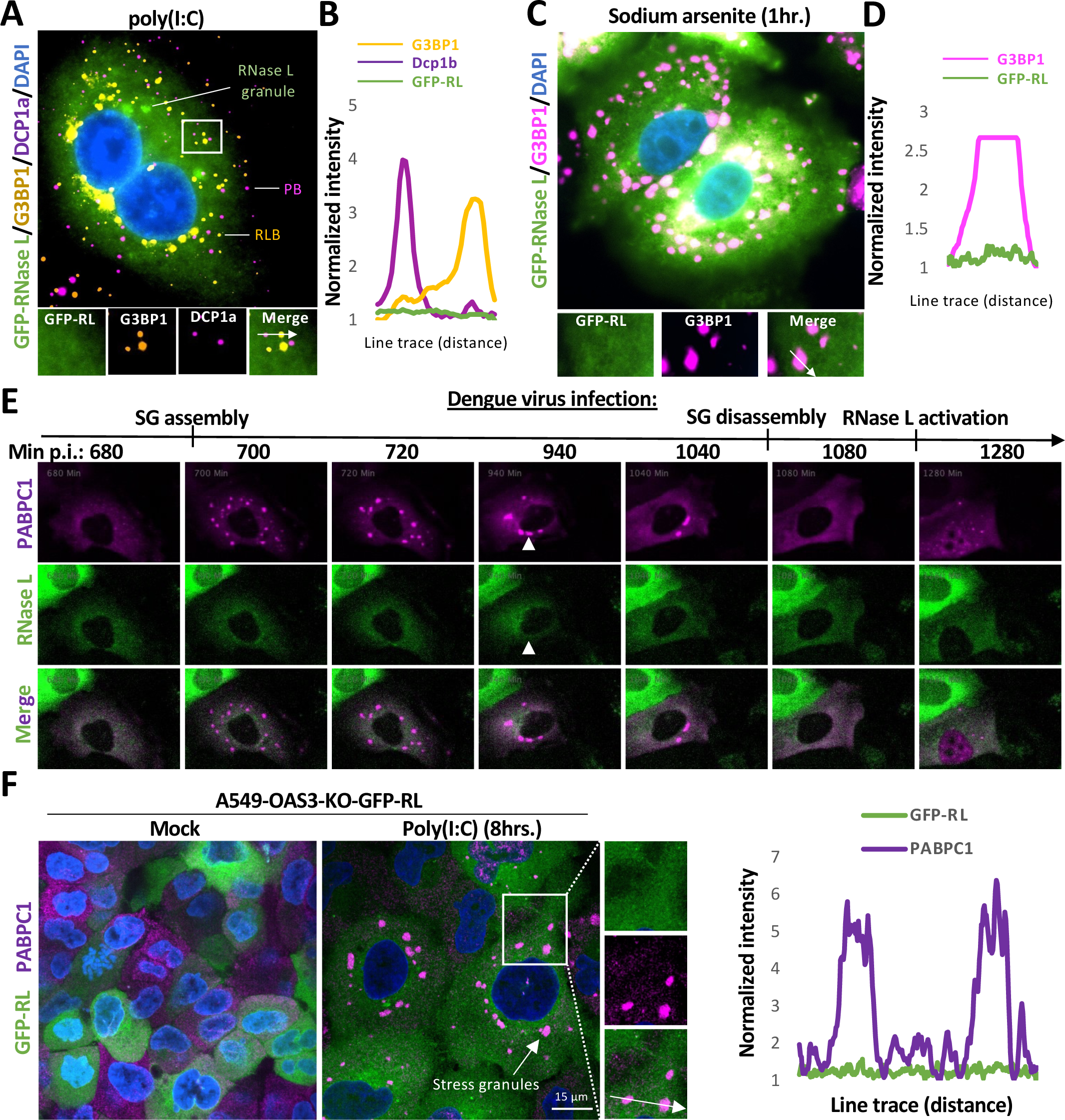
RNase L does not localize to P-bodies, RLBs, or stress granules. (A) Immunofluorescence staining for G3BP1 (RLB marker) and DCP1a (P-body marker) in RL-KO A549 cells expressing GFP-RNase L cells five hours post-lipofection of poly(I:C). (B) Line trace of SG, P-bodies, and RNase-L granule intensities. Intensities normalized to minimum intensity. (C) Immunofluorescence staining for G3BP1 (stress granule marker) in in RL-KO A549 cells expressing GFP-RNase L after 1 hour of 500 μM sodium arsenite treatment. D) Line trace through G3BP1 (SGs) and GFP-RNase L. (E) Live cell imaging of RL-KO A549 cells stably co-expressing both GFP-RNase L and mRuby2-PABPC1 infected with Dengue virus. (F) OAS3-KO cells expressing GFP-RL transfected with poly(I:C) and stained for PABPC1 (SG marker).

We next examined whether RNase L granules are stress granules, though we thought this was unlikely because RNase L activation inhibits the assembly of stress granules (Burke et al., 2019; Burke et al., 2020). We used three methods to generate stress granules in cells expressing GFP-RNase L. First, we treated cells expressing GFP-RNase L with sodium arsenite. Immunofluorescence assays for G3BP1 (stress granule marker) revealed that GFP-RNase L did not enrich in sodium arsenite-induced stress granules (Fig. 4C,D). Second, we infected cells expressing GFP-RNase L with dengue virus, which can occasionally induce stress granules prior to RNase L activation (Fig. 4E). Importantly, we did not observe RNase L enrich in dengue virus-induced stress granules (Fig. 4E and Movie 8). Lastly, we transfected OAS3-KO cells expressing GFP-RNase L with poly(I:C) and stained for PABPC1 (stress granule marker). Knockout of OAS3 phenocopied RNase L-KO cells, whereby PABPC1 remained in the cytoplasm and was incorporated into stress granules (Fig. 4F). However, GFP-RNase L did not enrich in dsRNA-induced stress granules in OAS3-KO cells (Fig. 4F). These data indicate that RNase L does not enrich in stress granules, contrary to previous reports (Reineke and Llyod, 2015; Paget et al., 2023).

### RNase L and OAS3 concentrate at double-stranded RNA foci

Recently, dsRNA-binding proteins such as PKR and ADAR1 were shown to concentrate in cytoplasmic condensates enriched for double-stranded RNA, termed double-stranded RNA-induced foci (dRIFs) (Zappa et al., 2021; Corbet et al., 2022). Because OAS3 binds dsRNA, we hypothesized that OAS3 could localize to dRIFs and lead to the concentration of RNase L at dRIFs. To determine if OAS3 localizes to dRIFs, we initially examined OAS3 localization via immunofluorescence assays in A549 cells following poly(I:C) lipofection. However, we were unsuccessful in detecting OAS3-specific immunofluorescence (Fig. S4A). Therefore, we tagged OAS3 on the N-terminus with mRuby-2 (Fig. S4B). We observed that OAS3 diffusely localized in the cytoplasm under mock transfection conditions (Fig. 5A).

**Figure 5.**
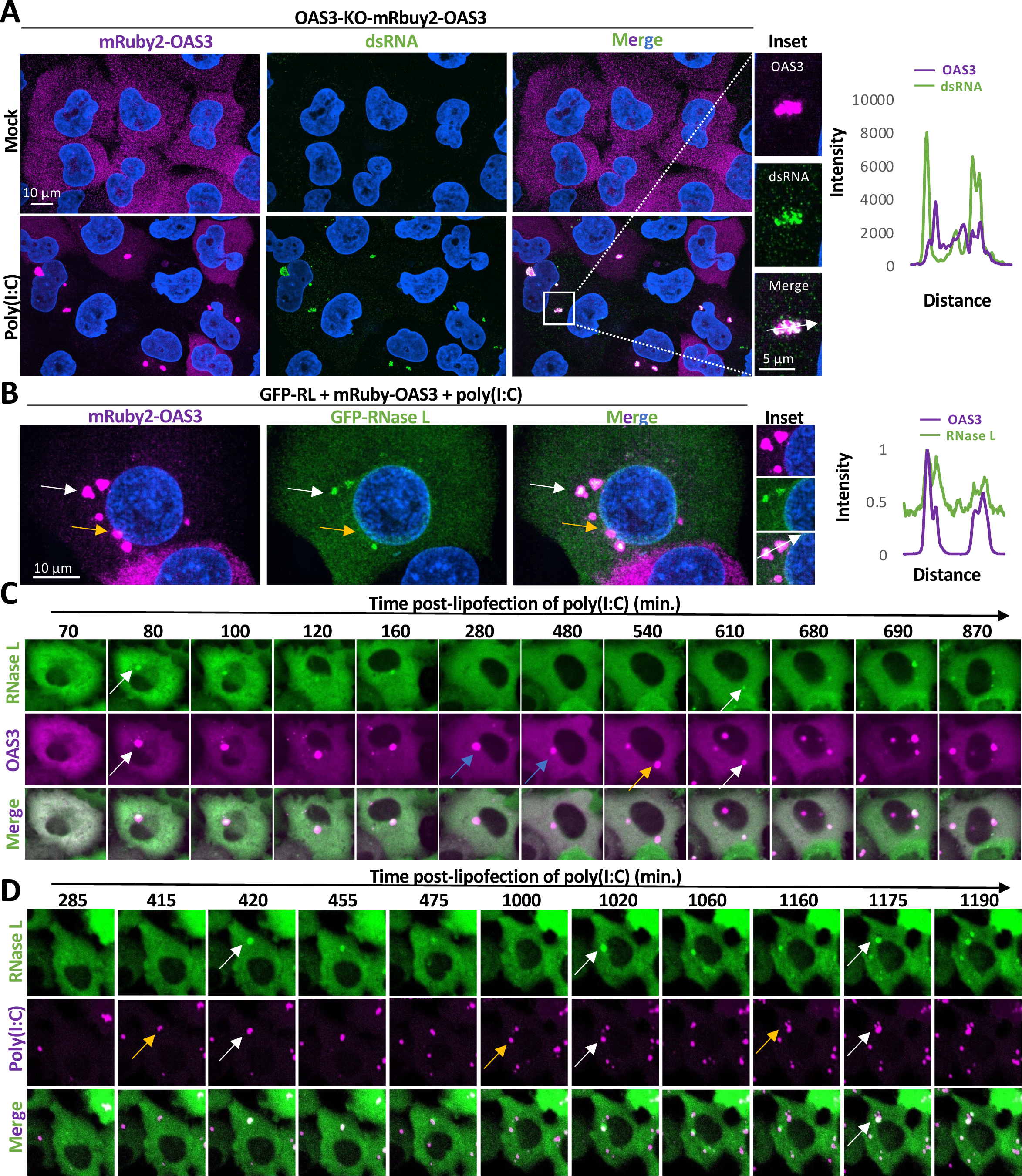
OAS3 and RNase L concentrate at double-stranded RNA-induced foci. (A) Fluorescence microscopy for mRuby-2-OAS3 and immunofluorescence staining for dsRNA in OAS3-KO A549 cells with or without lipofection of poly(I:C). Graph (right) shows intensity of line trace from inset. (B) Fluorescence microscopy of mRuby2-OAS3 A549 cells stably expressing GFP-RL transfected with poly(I:C). Graph shows intensity of line trace from inset. (C) Live cell imaging of RNase L-KO cells expressing GFP-RNase L and mRuby2-OAS3 following lipofection with poly(I:C). White arrows indicate co-incidental incorporation of GFP-RNase L and mRuby2-OAS3 into a dRIF. Blue arrows demarcate dRIFs that contain OAS3 (blue arrows) but never contain GFP-RNase L. Yellow arrows demarcate dRIFs that contain OAS3 but not RNase L, but then later incorporate GFP-RNase L (white arrow). (D) Live cell imaging of RNase L-KO cells expressing GFP-RNase L following lipofection with Rhodamine-labeled poly(I:C). Yellow arrows demarcate dRIFs that have not concentrated RNase L, but do concentrate RNase L at later times (white arrow).

Importantly, we observed that OAS3 concentrated into large granule structures that co-stained for dsRNA upon lipofection of poly(I:C) (Fig. 5A). Notably, mRuby2-OAS3 did not enrich in sodium arsenite-induced stress granules (Fig. S4C) nor RNase L-induced bodies (RLBs) (Fig. S4D,E). Moreover, mRuby2-OAS3 was functional because it rescued RNase L activation in OAS3-KO cells based on the observation that OAS3-KO cells expressing mRuby2-OAS3 displayed nuclear PABPC1 translocation and RLB assembly in response to poly(I:C) lipofection (Fig. S4F), which are markers of RNase L activation (Burke et al., 2019). In contrast, the parental OAS3-KO cells typically assembled stress granules and lacked nuclear PABPC1 following poly(I:C) lipofection (Fig. S4F), which phenocopies RNase L-KO cells (Burke et al., 2019). These data indicate that OAS3 concentrates at dRIFs.

To determine if OAS3 and RNase L co-localize at dRIFs, we generated cells that stably express both GFP-RNase L and mRuby2-OAS3. Following poly(I:C) lipofection, we observed that OAS3 and RNase L-co-localized into higher-order structures consistent with dRIFs (Fig. 5B, white arrow). Immunofluorescence assays demonstrated that GFP-RNase L co-localizes with PKR, phosphorylated PKR (p-PKR), and ADAR1 following poly(I:C) lipofection (Fig. S5A,B,C), which are known to localize to dRIFs (Zappa et al. 2021; Corbet et al., 2022). These data demonstrate that RNase L and OAS3 concentrate at dRIFs during the antiviral response.

Because some OAS3-positive dRIFs did not enriched for RNase L (Fig. 5B, yellow arrow), we hypothesized that RNase L transiently concentrates at OAS3-positive dRIFs. Indeed, live cell imaging of cells expressing GFP-RNase L and mRuby2-OAS3 following poly(I:C) lipofection demonstrated that RNase L typically concentrates at pre-formed OAS3-positive dRIFs but then dissipates over time (Fig. 5C and Movie 9). Similarly, we observed that RNase L transiently concentrates at pre-formed dRIFs following lipofection of rhodamine-labeled poly(I:C) (Fig. 5D and Movie10). These data indicate that RNase L episodically concentrates at OAS3-positive dRIFs.

### OAS3/RNase L-positive dRIFs assemble during viral infection

To determine if OAS3 and RNase L localize to dRIFs during viral infection, we infected A549 cells expressing GFP-RNase L and mRuby2-OAS3 with Zika virus (ZIKV) or Dengue virus serotype 2 (DENV2), which both activate RNase L (Whelan et al., 2019). We stained for viral RNA via smFISH. We observed that OAS3 and RNase L co-localized in higher-order assemblies near the ZIKV replication organelle (Fig. 6A). Moreover, live-cell imaging confirmed GFP-RNase L granules formed during dengue virus infection (Fig. 6B and Movie 11). Immunofluorescence staining for dsRNA and p-PKR showed that RNase L co-localized with viral dsRNA and p-PKR during dengue virus infection (Fig. 6C, white arrows). Notably, we also observed dRIFs that contained p-PKR and dsRNA but lacked RNase L (Fig. 6C, yellow arrow), indicating that RNase L transiently interacts with dRIFs and/or dRIFs are heterogenous in their composition. These data demonstrate that dRIFs assemble during Zika virus and dengue virus infection, and that RNase L and OAS3 concentrate at dRIFs.

**Figure 6.**
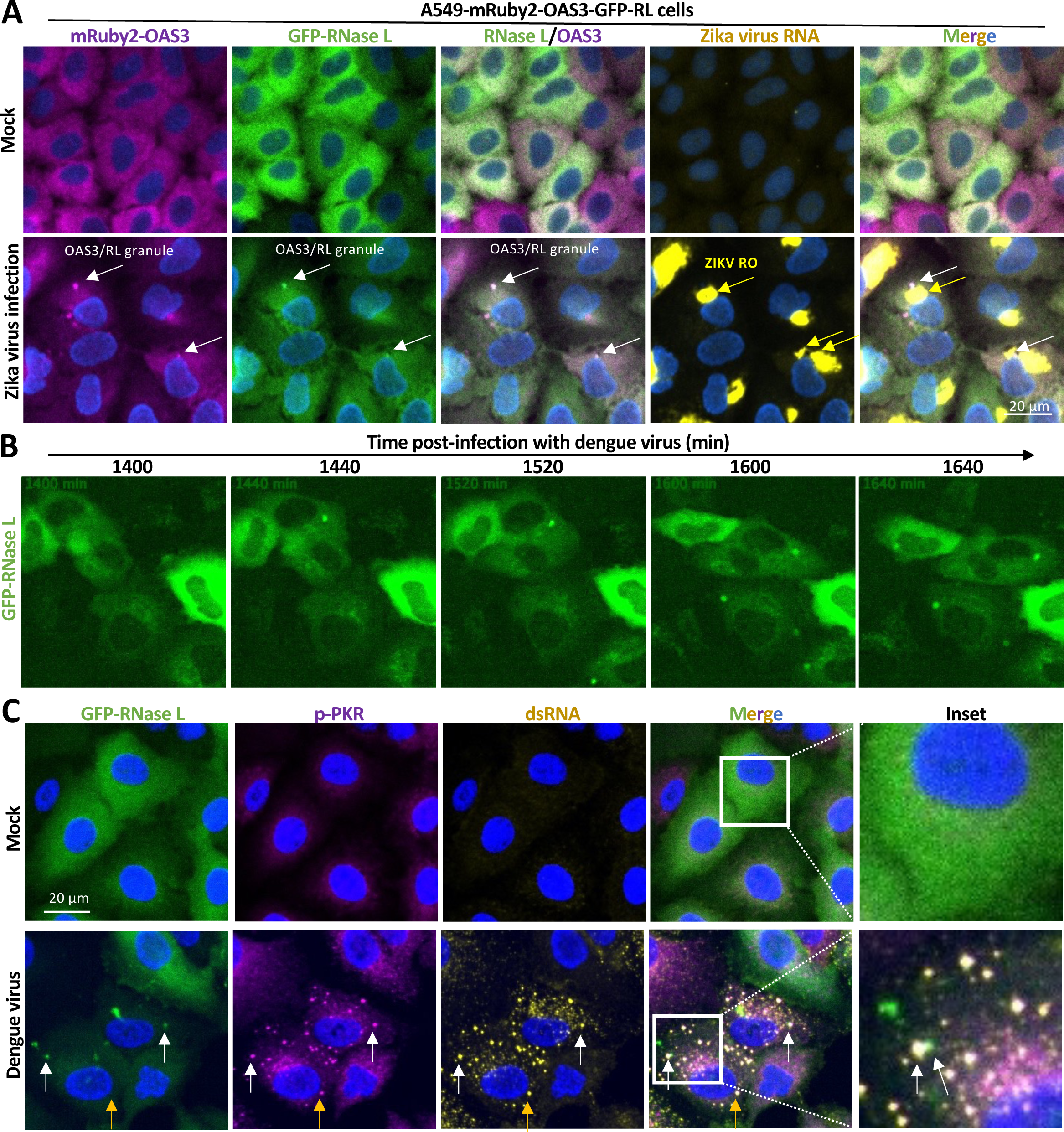
OAS3- and RNase L-positive dRIF assemblies form during viral infection. (A) smFISH/fluorescence microscopy of RNase L-KO A549 cells expressing GFP-RNase L and mRuby2-OAS3 following infection with Zika virus. White arrows indicate dRIFs while yellow arrows indicate zika virus replication factories. (B) Live cell imaging of RL-KO A549 cells expressing GFP-RNase L following dengue virus infection. (C) RNase L-KO cells expressing GFP-RNase L mock-infected and infected with dengue virus. White arrows indicate dRIFs that contain dsRNA, p-PKR, and GFP-RNase L. Yellow arrows demarcate dRIFs that contain dsRNA and p-PKR but not GFP-RNase L.

### RNase L and OAS3 dynamically interact with dRIFs

Our above data suggest that RNase L activates at dRIFs. One prediction of this model would be that RNase L dynamically interacts with dRIFs, allowing for exchange between the cytoplasm and dRIFs. To test this model, we examined whether GFP-RNase L displays fluorescence recovery after photobleaching (FRAP). Consistent with this hypothesis, we observed that GFP-RNase L readily recovers in dRIFs following photobleaching at early times post-lipofection of poly(I:C) (Fig 7A,B). Notably, this is when RNase L incorporation in dRIFs is transient (Fig. 2C). However, at later times post-poly(I:C) when RNase L becomes more stable in dRIFs (Fig. 2D,E), GFP-RNase L typically did not readily FRAP (Fig. 7B). These data indicate that RNase L readily exchanges between dRIFs and the cytoplasm at early times post-poly(I:C), but not at late time post-poly(I:C). Similar FRAP analyses of mRuby2-OAS3 revealed that OAS3 did not readily recover after photobleaching (Fig. 7C,D). This suggests that OAS3 stably associates with dRIFs or that dsRNA bound to OAS3 does not readily exchange with the cytoplasm. Combined, these data indicate that OAS3 concentrates in dRIFs by stably binding dsRNA, which in turn leads to the concentration of activated RNase L at OAS3-positive dRIFs (Fig. 7E).

**Figure 7.**
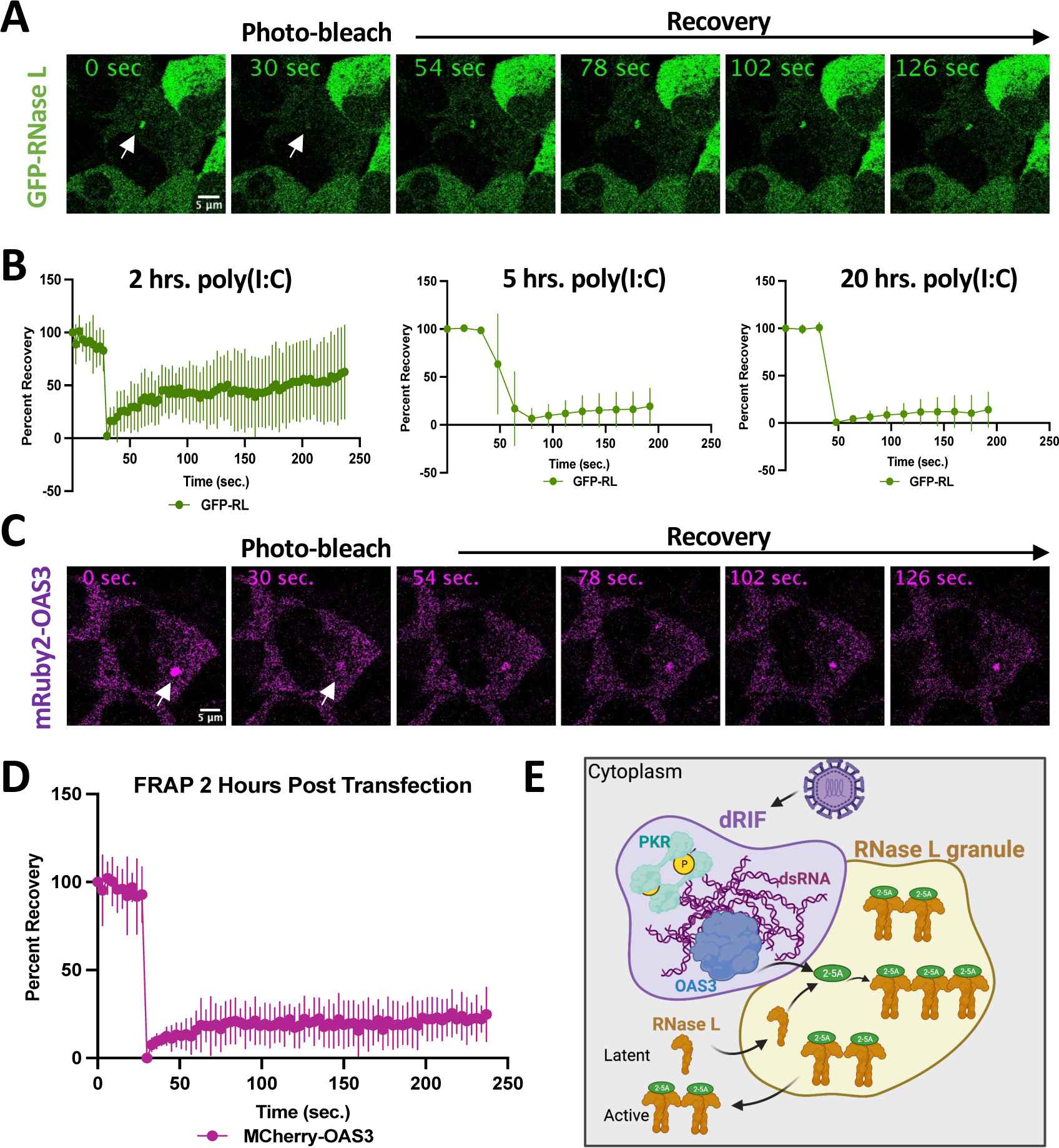
RNase L dynamically interacts with OAS3-positive dRIFs. (A-B) Fluorescence recovery after photobleaching (FRAP) analysis of RL-KO A549 cells expressing GFP-RNase L post-lipofection of poly(I:C). (A) Representative image of FRAP for GFP-RNase L two hours post-lipofection of poly(I:C).]. (B) Quantification of the average FRAP of GFP-RNase L in dRIFs 2-, 5-, and 20-hours post-poly(I:C) lipofection. (C-D) Similar to (A-B) but for mRuby2-OAS3 at two hours post-lipofection of poly(I:C). (D) Analysis of percent recovery of MRuby2-OAS3 in dRIFs after being photo-bleached. Analysis done for 2 hours post-poly(I:C) lipofection. (E) Model describing the interaction of RNase L with OAS3 at dRIFs.

## DISCUSSION

In this article, we examined the sub-cellular localization of RNase L and OAS3 during the antiviral response. We observed that OAS3 and RNase L are diffusely localized in the cytoplasm during homeostatic conditions but concentrate at double-stranded RNA-induced foci (dRIFs) in response to dsRNA or viral infection (Fig. 1,2,4,5,6). The concentration of RNase L at dRIFs coincides with RNase L activation (Fig. 3A,B) and is dependent on OAS3-mediated activation of RNase L (Fig. 3C,D). Moreover, activated (dimerized/oligomerized) RNase L enriches in dRIFs (Fig. 3G), and RNase L can readily exchange with the cytoplasm (Fig. 7A,B). Combined, these data support a model of OAS3/RNase L activation whereby dRIF assembly concentrates OAS3 and dsRNA, which supports efficient RNase L activation by concentrating RNase L to promote dimerization and oligomerization (Fig. 7E).

OAS proteins and RNase L were reported to co-localize with G3BP1 in stress granules driven by G3BP1 overexpression (Reineke and Lloyd, 2015). Moreover, two more recent studies reported that OAS and RNase L co-localize to stress granules in response to dsRNA (Manivannan et al., 2020; Paget et al., 2023). In contrast to these studies, our data strongly indicate that neither RNase L nor OAS3 co-localize with G3BP1 in stress granules that assemble in response to dsRNA, viral infection, or sodium arsenite (Fig. 4A, S4C,D). Moreover, neither OAS3 nor RNase L enrich in RNase L-induced bodies (RLBs) (Fig. 4A, S4E,F), which also concentrate G3BP1 following RNase L activation (Burke et al., 2020). Notably, while PKR was also previously shown to co-localize with G3BP1 in stress granules (Reineke and Lloyd, 2015; Manivannan et al., 2020; Paget et al., 2023), two recent studies conclusively demonstrated that PKR does not co-localize with G3BP1 in stress granules or RLBs (Zappa et al., 2021; Corbet et al., 2022). These studies combined with our findings herein question the long-standing notion that antiviral proteins (e.g., PKR, OAS3, and RNase L) localize to G3BP1 RNP complexes (stress granules and RLBs) to regulate antiviral signaling. Because G3BP1 and/or stress granules are reported to display antiviral activities (Onomoto et al., 2012; Yoo et al., 2015; Bidet et al., 2014; Galan et al., 2017), these observations warrant further examination into the mechanism by which G3BP1 and/or stress granules antagonize viral replication.

While RNase L and OAS3 do not localize to stress granules, they do localize to double-stranded RNA-induced foci (dRIF) (Fig. 5). Recently, PKR was shown to localize to dRIFs (Zappa et al., 2021; Corbet et al., 2022). Thus, antiviral protein localization at dRIFs appears to be a common feature of activating the antiviral response. How dRIFs assemble is unknown. One possibility is that dsRNA may be prone to condensation at high concentrations, which would be expected to occur in the viral replication organelle. However, we observed that dRIFs assembled outside the Zika virus and dengue virus replication organelle (Fig. 6A,B,C). This observation suggests that viral dsRNA leaks out of viral replication organelles, leading to dRIF assembly in the cytoplasm. Whether cellular factors in the cytoplasm promote the condensation of dsRNA into dRIFs is unknown. However, because many of the dsRNA-binding proteins (i.e., PKR, OAS3, ADAR1) that localize to dRIFs oligomerize upon activation, it is possible that oligomerization of dsRNA-binding proteins could promote multivalent interactions with dsRNA to seed dRIF assembly. Because concentration of antiviral proteins at dRIF assemblies could also promote antiviral protein dimerization/oligomerization, such as RNase L and PKR, the assembly of dRIFs could be a feed forward mechanism to rapidly potentiate antiviral signaling. Future studies will explore the mechanisms by which dRIFs assemble.

Similar to PKR and ADAR1, OAS3 likely concentrates at dRIFs via its ability to bind dsRNA-binding. Consistent with this, OAS3 interactions with dRIFs were not highly dynamic (Fig. 7), indicating that dsRNA binding affinity largely drives OAS3-dRIF interactions. However, an unresolved issue is understanding the interactions that lead to RNase L concentration at dRIFs. One possibility is that high concentrations of long 2-5A oligomers being synthesized by OAS3 could lead to a phase transition that localizes 2-5A near dRIFs, which then concentrates activated RNase L at dRIFs. Another possibility is that the dimerization/oligomerization of RNase L at dRIFs condenses RNase L into condensate-like state. Future work will focus on elucidating the molecule interactions of RNase L that lead to concentration at dRIFs.

An important observation we made is that the concentration of RNase L at dRIFs is transient during early times post-lipofection of poly(I:C) (Fig. 2C,D). This suggests that RNase L concentration in dRIFs is dynamic during this time, which is consistent with our FRAP analyses (Fig. 7A,B). However, at later times post poly(I:C), we observed that RNase L more stably associated with dRIFs (Figs. 2C,D, 7B), which suggests that RNase L exchange between dRIFs and the cytoplasm is reduced at later times post-poly(I:C). Notably, RNase L rapidly degrades cytoplasmic RNA following poly(I:C) lipofection (Burke et al., 2019; Cusic et al., 2023). Thus, we propose that a reduction in the interactions between RNase L and RNA in the cytoplasm at late times post-poly(I:C) could lead to an increase in the dwell time of RNase L molecules in dRIFs. Future work will explore the possibility that this is a mechanism to store activated RNase L and/or an autofeedback loop to reduce RNase L activity in the cytoplasm via sequestration to regulate cellular RNA decay.

Our observation that dRIFs containing dsRNA, p-PKR, OAS3, and RNase L form during Zika virus and dengue virus infection is significant because limited evidence previously existed that dRIFs form during viral infection, as Zappa et al. only observed PKR foci in a fraction of cells infected with measles virus (Zappa et al., 2021). Similarly, we observed dRIFs in only a small fraction of cells infected with either dengue virus or Zika virus. These observations indicate that dRIFs are either transient in nature during viral infection and/or viruses actively prevent dRIF assembly to prevent activation of antiviral signaling pathways. Future studies will focus on how viruses potentially prevent the condensation of their dsRNA intermediates into dRIFs.

In conclusion, we showed that several antiviral proteins, including PKR, OAS3 and RNase L, concentrate in dRIF assemblies that form in the cytoplasm near viral replication organelles. Our data demonstrating that RNase L concentrates at dRIFs upon activation argues that dRIFs are a site of activation for antiviral signaling in response to dsRNA or viral infection. Because dRIFs were only recently uncovered and little is understood about their assembly and function, future work characterizing these antiviral biological condensates will undoubtedly reveal fundamental insight into cellular surveillance of viral dsRNA and the initiation of antiviral signaling cascades.

## MATERIALS AND METHODS

### Cell culture

Parental and RNase L-KO (RL-KO) A549 cell lines and HEK293T cell lines are described in Burke et al., 2019. The A549 OAS3-KO cell line was a gift from Dr. Susan Weiss (Li et al., 2016). Cells were maintained at 5% CO_2_ and 37 degrees Celsius in Dulbecco’s modified eagle’ medium (DMEM) supplemented with fetal bovine serum (FBS; 10% v/v) and penicillin/streptomycin (1% v/v). Cells were transfected with poly(I:C) HMW (InvivoGen: tlrl-pic) or rhodamine-labeled poly(I:C) (InvivoGen: tlrl-picr) using 3-μl of lipofectamine 2000 (Thermo Fisher Scientific) per 500-ng of poly(I:C).

### Plasmids

To generate the GFP-tagged RNase L, eGFP was amplified via PCR from pLJM-1-eGFP (Addgene: Plasmid #19319) using primers that overlapped with pLenti-EF1-Blast at the xho1 site on the 5’ end and overlapped with the 5’-end of the RNase L coding sequence on the 3’-end (Supplementary table 1) and NEBNext® High-Fidelity 2X PCR Master Mix (New England Biolabs: M0541S). RNase L coding sequence was amplified using a 3’-end primer that overlapped with pLenti-EF1-Blast at the xba1 site. The fragments were gel purified and then assembled into a plasmid using in-fusion reagents (Takara Bio). The mCherry11-RNase L and mCherry1-10-RNase L were made similarly with oligos in Supplementary table 1. mCherry11-RNase L was inserted into pLenti-ef1-blasticidinR vector, whereas mCherry1-10-RNase L was inserted into pLenti-PGK-PuromycinR plasmid (Addgene: Plasmid #19070). The mRuby-OAS3 CDS and mRuby2 were amplified via PCR and primers (Supplemental table 1) and inserted into pLenti-PGK-PuromycinR via Gibson assembly (New England Biolabs: E2611S). The pLenti-mRuby2-PABPC1 is described in Burke et al., 2020.

### Viral infections

Dengue virus stereotype 2 (NGC) and Zika virus (PB 81) stocks were generated in Vero cells and titer was determined via plaque assay on Vero cells. A549 cells were infected at an MOI=0.5 by incubating cells (12-well format) with 0.5-ml of FBS-free medium containing virus for two hours with periodic rocking. Normal growth medium was added after two hours, and cells were fixed twenty-four hours post-infection. All viral infections were performed in a BSL3 facility at The Herbert Wertheim UF Scripps Institute for Biomedical Innovation & Technology using and under BSL3 biosafety protocols.

### Generation of GFP-RL and GFP cell lines

HEK293T cells (T-25 flask; 80% confluent) were co-transfected with 2.4-ug of lentiviral transfer plasmid, 0.8-ug of pVSV-G, 0.8-ug of pRSV-Rev, and 1.4-ug of pMDLg-pRRE using 20-ul of lipofectamine 2000. Media was collected at twenty-four-and forty-eight-hours post-transfection and filter-sterilized with a 0.45-um filter. To transduce A549 cell lines, A549 cells were incubated for 1 hour with 1-ml of lentivirus stock. Normal medium was then added to the flask and incubated for twenty-four hours. Medium was removed 24 hours post-transduction and replaced with selective growth medium containing 2-ug/ml of puromycin or blasicidin (Sigma-Aldrich). Selective medium was changed every three days. After one-week, selective medium was replaced with normal growth medium. Expression of proteins was confirmed via immunoblot analysis.

### Single-molecule fluorescent in situ hybridization (smFISH) and immunofluorescence assays

smFISH was performed following manufacturer’s protocol (https://biosearchassets.blob.core.windows.net/assets/bti_custom_stellaris_immunofluorescence_seq_protocol.pdf) and as described in Burke et al., 2019 and Burke et al., 2021. GAPDH probes labeled with Quasar 570 Dye were purchased from Stellaris (GAPDH: SMF-2026-1).

For immunofluorescence detection, cells were incubated with Rabbit polyclonal anti-PABP antibody (Abcam: ab21060) (1:1000), Mouse monoclonal anti-G3BP antibody (Abcam: ab56574) (1:1000), Recombinant Anti-RNase L antibody (Abcam: ab191392) (1:500), Mouse monoclonal RNase L Antibody (2E9) (Novus: NB100-351) (1:500), and Rabbit polyclonal OAS3 Antibody (Novus: NBP2-94081) (1:500) primary antibodies for four hours, washed three times, and then incubated with Goat Anti-Rabbit IgG H&L (Alexa Fluor® 647) (Abcam: ab150079) and Goat Anti-Mouse IgG H&L (FITC) (Abcam; ab97022) at 1:1000 for two hours. After three washes, cells were fixed and then smFISH protocol was performed.

### Microscopy and Image Analysis

Microscopy was performed as described in Burke et al., 2021. Briefly, cover slips were mounted on slides with VECTASHIELD Antifade Mounting Medium with DAPI (Vector Laboratories; H-1200). A Nikon Eclipse Ti2 equipped with a Yokogawa CSU-W1 Spinning Disk Confocal, 100x, 1.45 NA oil objective was used for imaging with a Nikon elements software. Tokai ThermoBox - Enclosure/Stage Top Incubation (Blackout) was used to incubate the cells for live cell imaging.

Deconvoluted images were processed using ImageJ with FIJI plugin. Z-planes were stacked, and minimum and maximum display values were set in ImageJ for each channel to properly view fluorescence. Fluorescent intensities and smFISH foci were quantified in ImageJ. Independent replicates were performed to confirm results.

### Fluorescent Recovery After Photo-Bleaching (FRAP) Analysis

FRAP was performed on a FV3000 confocal microscope with Abbelight 180 3D dStorm TIRF module, Tokai Hit STX environmental chamber, and Olympus software. Cells were seeded in a 4-chamber 35 mm glass bottom dish. A549 RL-KO GFP-RL MRuby2-OAS3 cells were transfected with 250 ng poly(I:C). Cells transfected for 2, 5, and 20 hours at 37 degrees Celsius before photobleaching. Mock wells were left as needed. Cells were placed in the environmental chamber with 5.0% CO_2_ and heat placed at 37 degrees Celsius for the duration of imaging. Lasers 488 and 570 were used for imaging and laser 405 (DAPI) was used for photobleaching. The 100X Oil High Resolution TIRF objective N.A. 1.5 was used to image. To photo-bleach, 405 laser intensity was set to 7% for a duration of 7 seconds to bleach to approximately cytoplasmic background of the cell. Recovery of the granule was monitored for 4 minutes. If the granule was still present after photo-bleaching, a second round of photobleaching was performed. To analyze the recovery, the first measurement pre-photobleaching was used for each granule to normalize all measurements to one.

### Crude Nuclear/Cytoplasmic Fractionation

Crude nuclear-cytoplasmic fractionation was performed as described in Burke et al., 2017. Parental WT and RL-KO GFP-RL A549 cells were seeded in a 12 well plate, 2 wells of each. Twenty-four hours later, one well of each was transfected with 500 ng poly(I:C). After six hours, cells were trypsonized, resuspended in 1 ml DMEM, and placed in 1.5 ml microcentrifuge tubes. Cells were pelleted at 500 × g for 5 minutes. After washing with phosphate buffered solution, the cell pellet was resuspended in 100 ul CSK buffer. 50 ul of resuspension was placed in fresh microcentrifuge tube and nuclei were pelleted at 5000xg for 10 min. 50 ul of 2X SDS Lysis buffer was placed in remaining 50 ul of resuspended pellet to be used as the whole cell lysate. The same process was repeated after chilling on ice for 10 minutes with repelleted aliquot to serve as the cytoplasmic lysate. Remaining pellet was washed with CSK buffer, aspirated and resuspended in 50 ul of CSK buffer and 50 UL of 2X SDS Lysis buffer as the nuclear fractionation.

### Sodium Arsenite Treatment

Sodium arsenite was suspended to a 100x 50mM stock in RNase free water. Final concentration of 500 um was used in 1 ml of media. Sodium arsenite treatment was left at 37 degrees Celsius for 1 hour before media was removed. Cells were then fixed with 4% PFA and permeabilized with 0.1% Triton-X.

## ACKNOWLEDGMENTS

The authors thank Dr. Roy Parker for providing reagents and valuable commentary. We thank Dr. Susan Weiss for the OAS3-KO A549 cell line. We thank Dr. Hyeryun Choe and Dr. Lizhou Zhang for providing Zika virus and dengue virus stocks.

## Author contributions

J.M.B. conceived the project. J.M.B. and R.C. performed experiments, analyzed data, and generated figures. R.C. and J.M.B wrote the manuscript.

## SUPPLEMENTAL FIGURE LEGENDS

**Figure S1. Immunofluorescence assay for RNase L. (**A-B) Immunofluorescence microscopy for RNase L using commercial antibodies in parental WT and RL-KO A549 cells. (A) Abcam RNase L antibody used for immunofluorescence assay. (B) Novus RNase L antibody used for immunofluorescence assay.

**Figure S2. RNase L is expressed in the cytoplasm during homeostatic conditions independent of cell cycle. (**A-B) Live cell imaging of sub-confluent RL-KO GFP-RL cells. (A) Sub-confluent cells displaying cytoplasmic localization of RNase L (left). Same cells imaged again 24 hours later when cells had become confluent (right). B, cells imaged through multiple cell divisions displaying cytoplasmic RNase L localization.

**Figure S3. RNase L granules are OAS3-dependent.** OAS3-KO cells co-expressing mRuby2-PABPC1 and GFP-RNase L post poly(I:C) lipofection.

**Figure S4. Immunofluorescence assays for OAS3. (**A) IF using commercial antibody for OAS3 in indicated cell lines (B) Western blotting assay for OAS3 and *GAPDH* mRNA showing mRuby2-OAS3 at about 140 kb. (C) Immunofluorescence assay for PABPC1 (SG marker) and fluorescence microscopy for MRuby2-OAS3 and GFP-RNase L in indicated cell line both shown under mock and post Sodium arsenite treatment. (D) Immunofluorescence assay for PABPC1 (SG marker) and dsRNA in MRuby2-OAS3 expressing cells shown under mock and post poly(I:C) lipofection. (E) Line tracing for PABPC1 (RLB marker), dsRNA (dRIF marker), and OAS3 for inset in (D). (F) Immunofluorescence assay for PABPC1 (SG and RLB marker) in indicated cell lines under mock and post poly(I:C) conditions.

**Figure S5. RNase L colocalizes with RNA binding proteins found in dRIFs. (**A,B,C) Immunofluorescence assays for proteins known to concentrate in dRIFs in indicated GFP expressing cell lines under both mock and poly(I:C) conditions displaying dRIFs. (A) Immunofluorescence assay performed for PKR. (B) Immunofluorescence assay performed for P-PKR, cells transfected with Rhodamine poly(I:C). (C) Immunofluorescence performed for ADAR1 and dsRNA.

**Figure S6. FRAP analyses at late times post-lipofection of poly(I:C). (**A) GFP-RL positive granule photobleached post 20 hours poly(I:C) lipofection.

## SUPPLEMENTAL VIDEOS

**Movie 1. Related to Figure 1G.**

Live-cell imaging of GFP-RNase L (green) and mRuby2-PABPC1 (magenta) at indicated times post-lipofection of poly(I:C). The video shows that PABPC1 translocates to the nucleus following activation of RNase L and the formation of RNase L-induced bodies (RLBs). Note, GFP-RNase L granules can be observed.

**Movie 2. Related to Figure 1H.**

Live-cell imaging of GFP-RNase L (green) following RNase L activation. RNase L translocates to the nucleus, which is followed by cell death. Note, the video of this cells is an excerpt from a larger image (Movie 4).

**Movie 3. Related to Figure 2C.**

Live-cell imaging of GFP-RNase L (green) following RNase L activation. Video shows transient assembly of GFP-RNase L in an higher-order assembly.

**Movie 4. Related to Figure 2D.**

Live-cell imaging of GFP-RNase L (green) in A549 RNase L-KO cells expressing GFP-RNase L following lipofection of poly(I:C). This video starts at time=0 and runs to 260 min. post-lipofection of poly(I:C). Movie 5 is a continuation of this video (285-1200 min).

**Movie 5. Related to Figure 2D.**

Live-cell imaging of GFP-RNase L (green) in A549 RNase L-KO cells expressing GFP-RNase L following lipofection of poly(I:C). This video is a continuation of Movie 4 and starts at 285 min post-lipofection. The video shows that most cells assemble higher-order structures that contain GFP-RNase L. The assemblies are transient at earlier times, but become more stable at later times.

**Movie 6. Related to Figure 3A.**

Live-cell imaging of GFP-RNase L (green) and mRuby2-PABPC1 (magenta) at indicated times post-lipofection of poly(I:C) showing that RNase L granules typically form just prior to RLB assemble (small PABPC1+ foci).

**Movie 7. Related to Figure 3G.**

Live-cell imaging of GFP-RNase L (green) and split mCherry-RNase L reporters (magenta) at indicated times post-lipofection of poly(I:C).

**Movie 8. Related to Figure 4E.**

Live-cell imaging of GFP-RNase L (green) and mRuby2-PABPC1 (magenta) at indicated times post-infection with dengue virus. Video shows a cell that assemble PABPC1-positive stress granules, which do not enrich RNase L. After the stress granules disassemble, RNase L activation results in the assembly of small PABPC1 foci (RLBs) and translocation of PABP to the nucleus. RNase L does not enrich in RLBs.

**Movie 9. Related to Figure 5C.**

Live-cell imaging of GFP-RNase L (green) and mRuby2-OAS3 (magenta) at indicated times post-lipofection of poly(I:C).

**Movie 10. Related to Figure 5D**

Live-cell imaging of GFP-RNase L (green) and Rhodamine-labeled poly(I:C) (magenta) at indicated times post-lipofection of poly(I:C).

**Movie 11 Related to Figure 6D**

Live-cell imaging of GFP-RNase L (green) at indicated times post-infection with dengue virus.

