## Supplemental Figures for "OAS3 and RNase L integrate into higher-order antiviral condensates"

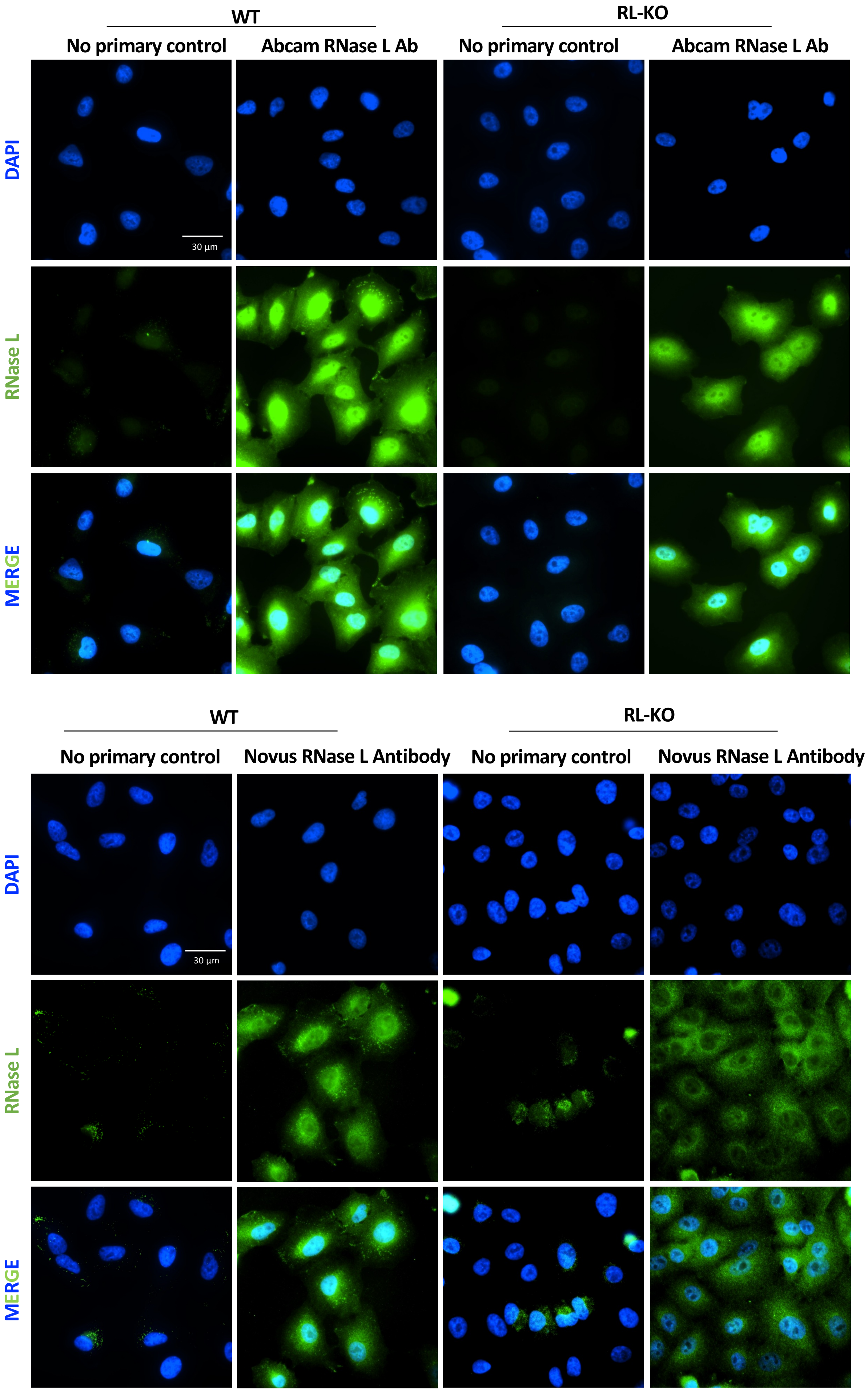
WT
RL-KO
No primary control
Abcam RNase L Ab
No primary control
Abcam RNase L Ab
DAPI
RNase L
MERGE
30 µm
WT
RL-KO
No primary control
Novus RNase L Antibody
No primary control
Novus RNase L Antibody
DAPI
RNase L
MERGE
30 µm

Fig. S1

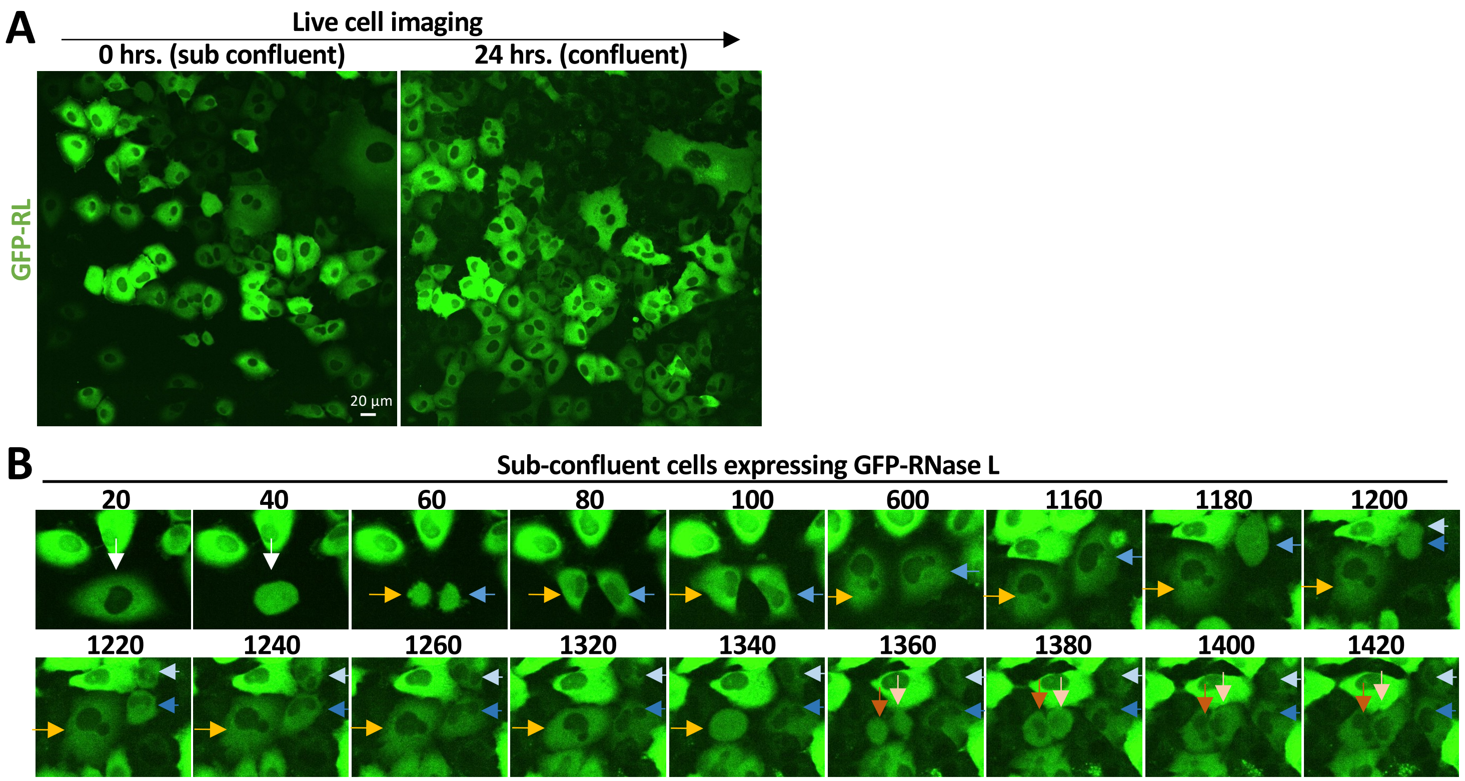
A
Live cell imaging
0 hrs. (sub confluent)
24 hrs. (confluent)
GFP-RL
20 µm
B
Sub-confluent cells expressing GFP-RNase L
20
40
60
80
100
600
1160
1180
1200
1220
1240
1260
1320
1340
1360
1380
1400
1420

Fig. S2

A549-OAS3-KO cells expressing GFP-RNase L post-lipofection of poly(I:C) (6 hrs.)

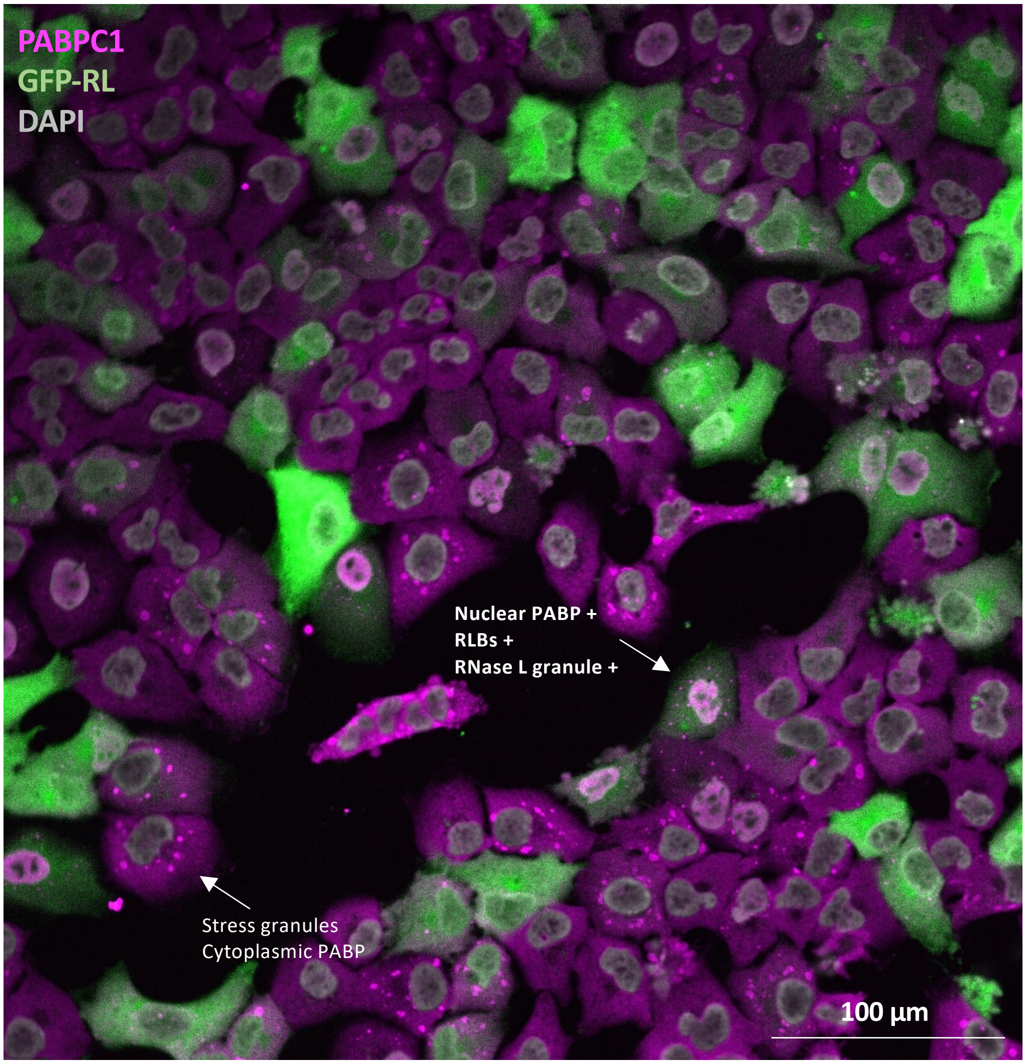


Fig. S3

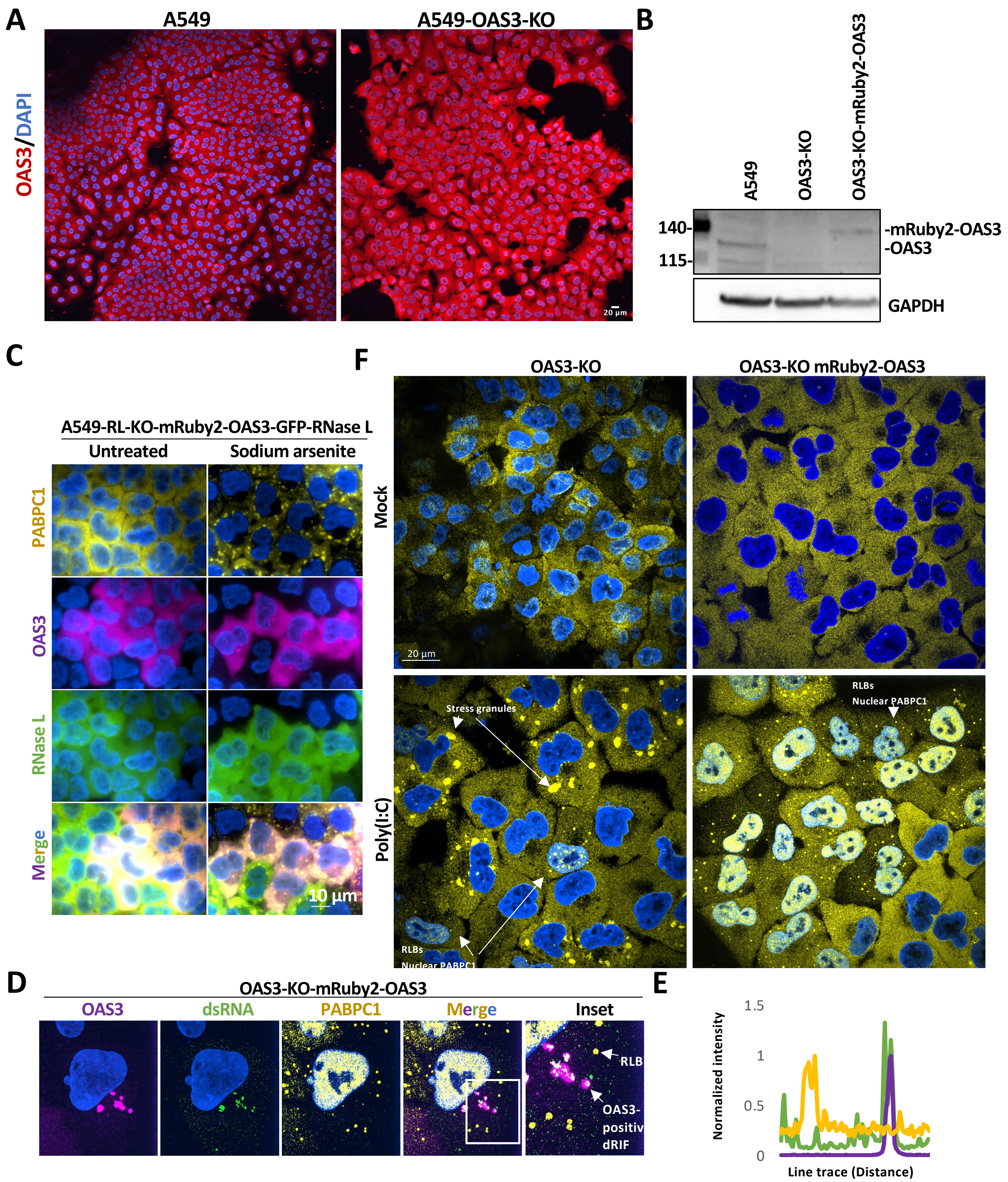
A
A549
A549-OAS3-KO
OAS3/DAPI
20 µm
B
A549
OAS3-KO
OAS3-KO-mRuby2-OAS3
140-
115-
-mRuby2-OAS3
-OAS3
GAPDH
C
A549-RL-KO-mRuby2-OAS3-GFP-RNase L
Untreated
Sodium arsenite
PABPC1
OAS3
RNase L
Merge
10 µm
F
OAS3-KO
OAS3-KO mRuby2-OAS3
Mock
Poly(I:C)
20 µm
Stress granules
RLBs
Nuclear PABPC1
D
OAS3-KO-mRuby2-OAS3
OAS3
dsRNA
PABPC1
Merge
Inset
RLB
OAS3-
positiv
dRIF
E
Normalized intensity
1.5
1
0.5
0
Line trace (Distance)

Fig. S4

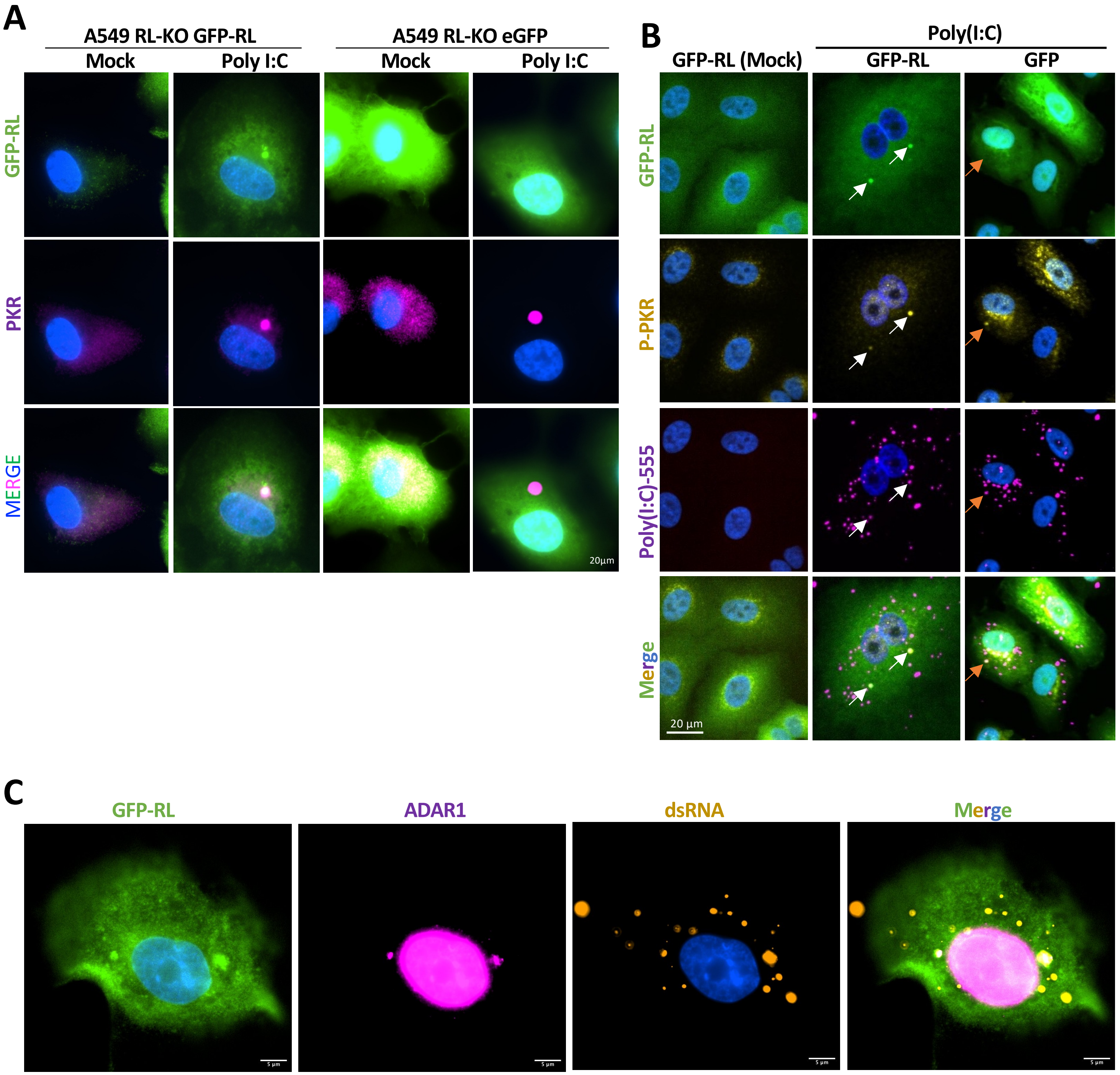
A
A549 RL-KO GFP-RL
Mock
Poly I:C
A549 RL-KO eGFP
Mock
Poly I:C
GFP-RL
PKR
MERGE
20µm
B
Poly(I:C)
GFP-RL (Mock)
GFP-RL
GFP
GFP-RL
P-PKR
Poly(I:C)-555
Merge
20 µm
C
GFP-RL
ADAR1
dsRNA
Merge

Fig. S5

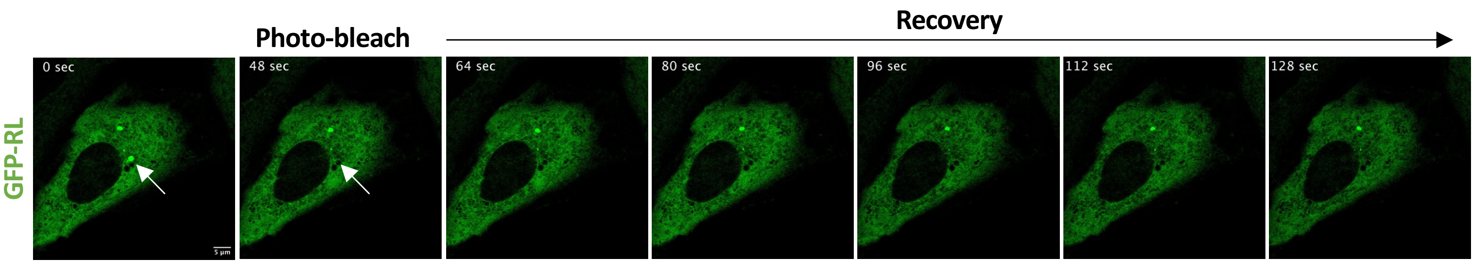
Photo-bleach
Recovery
GFP-RL
0 sec
48 sec
64 sec
80 sec
96 sec
112 sec
128 sec
5 µm

Fig. S6

**SUPPLEMENTAL FIGURE LEGENDS**

**Figure S1. Immunofluorescence assay for RNase L. (**A-B) Immunofluorescence microscopy for RNase L using commercial antibodies in parental WT and RL-KO A549 cells. (A) Abcam RNase L antibody used for immunofluorescence assay. (B) Novus RNase L antibody used for immunofluorescence assay.
